# Rat Hindlimb Amputation Model Shows Analgesia and Sexually Dimorphic Cold Hypersensitivity with Immediate Targeted Muscle Reinnervation

**DOI:** 10.1101/2025.03.06.641863

**Authors:** Jose Lucas Zepeda, Gabriella Mraz, Elizabeth Roth, Dorothee Weihrauch, Gwendolyn Hoben

**Author notes:** **Correspondence to:** Gwendolyn Hoben, MD, PhD, Department of Plastic Surgery, Medical College of Wisconsin, Gwendolyn Hoben. **Emails:** Jose Lucas Zepeda; Gabriella Mraz; Elizabeth Roth; Dorothee Weihrauch.

## Abstract

**Introduction:** Targeted muscle reinnervation (TMR) has been used clinically to reduce pain in amputees but the mechanisms producing analgesia are unknown. Here we employ rat hindlimb amputation as an appropriate model to evaluate algogenic processes affected by TMR, performed at the time of amputation in subjects of both sex.

**Methods:** Adult male and female Sprague-Dawley rats were assigned to amputation-only or amputation + immediate TMR (iTMR). Reflexive pain behaviors (von Frey, pin, and cold hypersensitivity testing) and spontaneous pain behaviors (guarding, flinching, anhedonia from loss of sucrose preference) were assessed. Retrograde labeling distinguished sensory and motor neurons and identified nerve coaptation sites.

**Results:** iTMR reduced hyperalgesia and anhedonia that followed amputation. By 4 weeks, pin testing showed reduced pain responses in iTMR rats (25%) versus amputation-only (51%, p=0.01). Sucrose preference testing revealed reduced anhedonia in iTMR rats (76% vs. 43%, p=0.01). Response to cold diminished in male iTMR rats (49% vs. 100% in amputation-only males, p<0.001), but not in females. Histology showed neuroma formation in controls but not after iTMR. Both sensory and motor fibers entered the motor branch after iTMR.

**Conclusions:** These findings show iTMR prevents neuroma formation and provides analgesia after amputation and, while its effect on cold hypersensitivity is sexually dimorphic. This supports further investigation into TMR analgesic mechanisms, the utility of the hindlimb amputation model, and the need for sex-specific evaluations.

**Perspective:** Using a rat hindlimb amputation model to evaluate iTMR, we demonstrate its potential to mitigate neuropathic pain and neuroma formation and reveal sex-specific responses to cold hypersensitivity. These findings indicate the potential clinical utility of iTMR in improving pain outcomes, potentially providing more tailored approaches to pain management in amputees.

## 1. Introduction

Chronic pain affects over 70% of major limb amputees.^1-3^ Prior to targeted muscle reinnervation (TMR) the available interventions, from medication to mirror therapy only provided incremental relief.^4,5^ TMR is a surgical procedure initially developed to improve myoelectric prosthesis control by redirecting amputated nerves to new muscle targets.^6^ It has been found to be very effective in that intended function but has found greater utility in reducing or preventing amputation related pain. A multi-institutional cohort study by Valerio et al. showed that 49% of patients who underwent early TMR within 2 weeks of amputation were completely free of residual limb pain (RLP) and 45% were free of phantom limb pain (PLP). In contrast, only 19% and 21% of standard amputation (no TMR) patients were free of RLP and PLP, respectively. Moreover, the TMR patients required less opioids.^7^ findings have greatly increased the utilization of TMR in amputees solely for the purpose of pain prevention and/or treatment.^7-9^ The mechanism of analgesic action of TMR is not understood. When TMR is performed, the amputated nerves are coapted to motor branches proximal to the injury. Clinically, this is a counter-intuitive procedure as the large amputated nerves, often mixed motor-sensory nerves, are sewn to much smaller motor-only branches. The amputated motor axons clearly re-innervate the new muscle and when stimulated, there is muscular contraction sufficient to create an EMG signal for controlling a prosthesis. However, the peripheral and central consequences of amputating a sensory nerve and connecting it to a motor branch are much less clear. Marasco et al. showed in patients that there was some sensory reinnervation to the skin from the amputated nerves when TMR was performed, but the lack of expected neuroma formation or neuropathic pain in patients that underwent TMR suggest an unexplained role of sensory reinnervation in influencing pain pathways.^10,11^

We previously examined TMR as an analgesic intervention in a spared nerve injury (SNI) model and demonstrated clear reduction in neuropathic pain behaviors.^12^ Senger et al also applied TMR in a rodent following tibial injury and showed improved analgesia compared to tibial nerve ligation.^13^ To move closer to the most common clinical setting of TMR, we sought to create a hindlimb amputation model for evaluating TMR analgesia. This would more accurately reflect the greater injury to the dorsal root ganglia (DRG) neurons as there would be many fewer healthy sensory neurons at the level of the injury, in contrast to SNI in which the sural nerve is retained.^12^ An amputation model may therefore provide a foundation for assessing rodent behaviors associated with neuropathic pain and evaluating the effects of interventions like TMR on pain modulation in the context of more severe injuries. Clinical work in amputation-related pain suffers from under-representation of female patients, and so inclusion of female animal subjects will also facilitate identification of sex-related differences in amputation-related pain.^3,7,8^

In the current study, we developed a rat hindlimb amputation model to investigate the effects of immediate targeted muscle reinnervation (iTMR) on amputation related pain. Both evoked and spontaneous pain behaviors were assessed. By advancing our understanding of the neurobiological effects of TMR with this animal model, we aim to contribute to the optimization of TMR procedures and improve clinical outcomes for amputees and those with more limited nerve injuries.

## 2. METHODS

### 2.1. Animal Operations

All animal experiments and surgical procedures were performed in compliance with the Association for Assessment and Accreditation of Laboratory Animal Care International and the Institutional Animal Care and Use Committee and National Institutes of Health guidelines. Male and female adult Sprague-Dawley rats (150-200g; Charles River Laboratories, Wilmington, MA) were obtained and used according to the protocols approved by the Medical College of Wisconsin. Animals of both sexes were randomly assigned to either of the following surgical interventions: hindlimb amputation only or hindlimb amputation and iTMR.

### 2.2. Surgical Procedures

Operative procedures were conducted aseptically in a dedicated operating space with an operating microscope (AmScope, Irvine, CA). Anesthesia was induced with inhaled 4% isoflurane (Phoenix, St. Joseph, MO), confirmed by absence of withdrawal response to toe-pinch, and maintained with 1.5% to 2.0% isoflurane. Rats were injected with 1 mg/kg meloxicam subcutaneously (VetONE, Down, Northern Ireland).

For the left hindlimb amputation procedure (**Fig. 1A,C**), a longitudinal incision splitting the gluteal and hamstring muscles on the lateral aspect of the thigh was made to expose the sciatic nerve just proximal to its trifurcation. The tibial, sural and peroneal nerves were ligated with 5-0 silk and transected. Proximally, the caudofemoralis muscle was divided to expose the branches of the sciatic to caudofemoralis, the branch to semimembranosus, and the branches of the biceps femoris. The branch to caudofemoralis was maintained so that there would not be nearby denervated muscle and the nerves used as recipients in the TMR operation were divided and the distal ends were flipped distally to prevent reinnervation. These nerves were transected to control for any loss of motor function related to the recipient nerves in the iTMR procedure. A separate circumferential incision 3cm distal to the knee joint was performed, the anterior and posterior compartment musculature was divided, and dissection proceeded proximally to expose 1cm of the tibia and fibula. These were divided sharply, hemostasis achieved, and the anterior and posterior compartment musculature were sewn together over the bone stumps. The posterior skin flap was brought anteriorly, and the anterior skin flap was shortened as needed to prevent the suture line from being on the most dependent aspect of the stump. The closure was then sealed with Vetbond™ (World Precision Instruments, Sarasota, Fla., USA).

**Figure 1:**
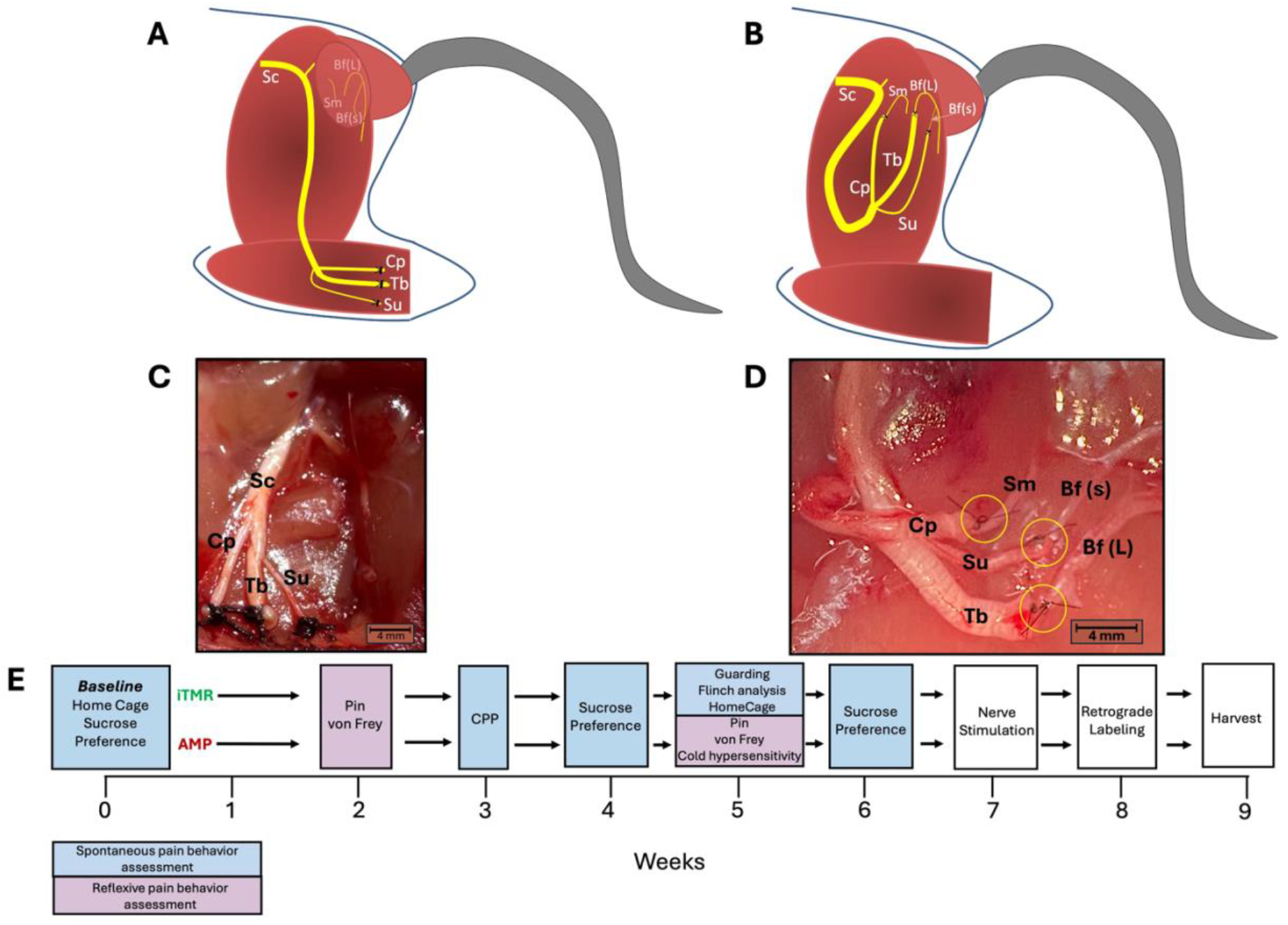
Surgical Procedure and Experimental Timeline for iTMR and AMP Models. (A) Schematic of below-the-knee amputation (AMP) on the left hind limb with ligation of the tibial (Tb), common peroneal (Cp), and sural (Su) nerves. The. (B) Schematic of targeted muscle reinnervation (iTMR) model, showing nerve coaptations: tibial nerve to the larger motor branch of the biceps femoris (Bf-L), common peroneal nerve to the motor branch of the semimembranosus (Sm), and sural nerve to the smaller motor branch of the biceps femoris (Bf-S), highlighting clinical size mismatches. (C) In vivo image of the AMP surgical model. (D) In vivo image of the iTMR surgical model. (E) Experimental timeline over 9 weeks, detailing assessments of spontaneous and reflexive pain behaviors, followed by nerve stimulation, retrograde labeling, and tissue harvest.

For animals receiving hindlimb amputation + iTMR (**Fig. 1B**), the same longitudinal incision was made to expose the sciatic nerve just proximal to its trifurcation. The tibial, sural, and common peroneal nerves were separated and dissected free from the main sciatic trunk. As above, the recipient nerves were dissected and divided distally. The ends of the tibial, sural, and peroneal nerves were coapted with 10-0 nylon suture (AROSurgical, Newport Beach, CA) to the recipient motor nerves, large branch to biceps femoris, small branch to biceps femoris, and the branch to semimembranosus, respectively (**Fig. 1D**). Following iTMR, the hindlimb amputation procedure was performed as described previously.

### 2.3. Postoperative Care

Animals were allowed to recover and were monitored for postoperative complications before being returned to the animal care facility. They were provided with food and water *ad libitum* and were monitored daily for signs of infection or stress. At the appropriate time point following intervention, animals were euthanized with 5% isoflurane and cervical spinal cord transection.

### 2.4. Retrograde Labeling and Tissue Collection

During anesthesia, the gluteal splitting incision was re-opened 8 weeks following the surgical procedures. The common peroneal nerve was carefully dissected. In the amputation only group, the nerve was divided just proximal to the ligature and in the iTMR group, the coaptation to the semimembranosus branch was carefully dissected and the branch to semimembranosus was divided. The cut nerve end was placed in a Vaseline well with 15μl of 4% Fluoro-gold (Fluorochrome, Denver, CO) solution for one hour. The nerve was then gently washed of excess dye, and the incision was re-closed in layers. One week later, the L5 DRG and the spinal cord were harvested, fixed in 4% paraformaldehyde followed by 30% sucrose for cryoprotection, and frozen in optimal cutting temperature (OCT) compound (VWR, Radnor, PA). The DRGs were then sectioned at a thickness of 20 μm and the cords at 50 μm. The number of fluoro-gold positive nuclei were counted on a Nikon Eclipse Ti2 microscope (Nikon Inc, Melville, NY). For DRGs, every 5th section was counted while the entirety of the lumbar ventral horn was analyzed. Each section was counted independently by 2 of the authors (GM, JLZ) and the counts averaged for analysis. A 40% correction factor was applied to account for double count errors. ^14,15^ The tibial and sural nerve ligated ends (amputation only) and coaptations (iTMR group) were also collected at this time and prepared for paraffin embedding.

### 2.5. Nerve Stimulation

Prior to dividing the common peroneal for retrograde labeling, the nerves were stimulated to assess reinnervation. The tibial, sural, and common peroneal nerves proximal to the original transection or coaptation, were electrically isolated from each other with nitrile squares. A nerve stimulator (Checkpoint Surgical, Cleveland, OH) producing individual pulses with 0.05 mA and 100 μs duration was used to stimulate each nerve separately. The presence or absence of muscle twitch in response to stimulation was recorded.^12^

### 2.6. Pain Behavior Measurements

Testing was performed with the rats placed individually in clear plastic enclosures (13 × 25 × 25 cm) on a ¼-in wire grid, unless otherwise stated. All the animals were habituated in the testing room for 1 hour. All behavior testing was performed by individuals blinded to the intervention group. All stimuli were applied to the amputated left central stump. Baseline behaviors were obtained pre-operatively (**Fig 1E**). At 2 and 5 weeks post-operatively, von Frey (mechanical hypersensitivity) and pin (hyperalgesia) responses were tested. Conditioned placed preference (CPP) with gabapentin was completed three weeks post-operatively. Sucrose preference testing was performed at baseline and 4 weeks post-intervention. Acetone response (representing cold hypersensitivity), guarding (spontaneous pain), flinching (spontaneous pain) and HomeCageScan were assessed 5 weeks post-operatively. Animals were excluded from testing if found sleeping, resting with closed eyes and stomachs on the mesh, or engaged in distracting behaviors such as grooming, eating, rearing, or walking.

#### 2.6.1. Evoked Pain Measures

Three distinct evoked pain measures were employed to assess nociceptive behavior. **von Frey** testing used a series of monofilaments (Performance Health, Warrenville, IL), numbered 2-9, which exerted varying forces corresponding to their number, with lower numbers producing lesser force and higher numbers producing greater force. Testing commenced with filament 5, applied to the center of the left hind paw. Responses such as lifting the stump, fluttering, licking, or the absence of movement were recorded. An up-down paradigm was employed to determine sensitivity, and the results were expressed as the 50% withdrawal threshold in grams.^16^ **Pin testing** involved touching the plantar skin with a 22G spinal needle (BD, Franklin Lakes, NJ) that elicit a withdrawal response without breaking the skin.

Responses were categorized into one of 3 types: hyperalgesic response (stump lift and held, flutter, or lick), normal withdrawal response (stump lift and immediate replacement back on the mesh), or no response.^17,18^ The hyperalgesic response has been identified as selectively indicting the affective unpleasantness element of pain as it alone is associated with place avoidance.^19^ The data were presented as a percentage of hyperalgesic responses across 10 test applications. The **acetone test** involved extruding

0.2 mL of acetone (EKI Chem, St. Joliet, IL) from a 25G blunt needle (BD) and applying it to the center of the stump, causing evaporative cooling. The frequency of hyperalgesic responses as described above was assessed over 5 trials.^20,21^

#### 2.6.2 Spontaneous Pain Measures

The **Sucrose Preference Test** was performed as an indicator of anhedonia. Animals were acclimated to two bottles, one containing 1% sucrose solution and the other water, over a 5-day period, with alternating bottle placement to prevent side preference. On the test day, after an 8-hour deprivation of food and water, sucrose consumption over 16 hours was recorded. Measurements were taken preoperatively and at 4 and 6 weeks postoperatively.^22,23^ The **Flinching Score** was used to quantify spontaneous pain by recording the number of times the animal raised its hind paw. After acclimating to the testing room for an hour, flinches were recorded during a two-minute observation period.^24^ The **Guarding Score** was assessed at 5-minute intervals over the course of one hour. The left hindlimb was rated on a scale from 0 to 2: a score of 0 indicated the stump rested on the cage floor (no spontaneous pain), a score of 1 was given if the stump touched the mesh but without distortion of the skin, indicating minimal pressure, and a score of 2 indicated full guarding, where the paw was elevated off the floor mesh (indicative of spontaneous pain behavior).^25^ The **Conditioned Place Preference (CPP)** test identifies if rats are experiencing neuropathic pain by determining if the established analgesic effects of gabapentin motivate the subject to prefer the chamber in which it was provided. This test was performed 3 weeks after surgical intervention using a 2-chamber setup separated by a removable wall, with distinct visual (either stripes or dots on the chamber walls) plus olfactory cues (either ChapStick mint or cherry) paired with gabapentin (Sigma-Aldrich, St.

Louis, MO) versus saline administration to assess conditioned preferences. The experiment spanned 6 consecutive days. On day 1, animals were acclimated to the testing suite and the CPP box for 30 minutes, with open access to both chambers. On day 2, preconditioning took place, during which animals had free access to both chambers for 15 minutes to assess baseline preferences. On days 3, 4, and 5, a biased drug-pairing procedure was conducted, during which animals received intraperitoneal (IP) injections of either saline or gabapentin (100 mg/kg), with each injection separated by 4 hours. Saline was paired with the striped chamber and berry scent in the morning, while gabapentin was paired with the polka-dot chamber and vanilla scent in the afternoon. Forty-five minutes post-injection, based on previous measure of the onset of gabapentin analgesia, animals were placed in the corresponding chamber for 15 minutes without access to the other side.^26^ On day 6, postconditioning testing was performed,during which animals were given open access to both chambers for 15 minutes to evaluate the acquisition of CPP. The difference score was calculated by subtracting the time spent in the drug-paired chamber after conditioning from the time spent in the same chamber before conditioning.^27^ **HomeCageScan (HCS)** is an automated video-based behavioral scoring system to assess spontaneous and ongoing pain behaviors in rats within their home cage environment (Clever Systems Inc., Reston, VA). Rats were individually housed in clear plastic cages (42 cm × 18 cm × 24 cm) for the duration of the analysis, with food and water provided ad libitum. All video recording was performed during the dark phase, when rats are typically most active, using infrared backlighting to allow for uninterrupted observation. The HCS system, operated using Clever Systems software version 3.00, exported behavioral data in 1-minute intervals and subsequently converted it into the proportion of time spent on each behavior.^28^ Prior to recording, animals were acclimated to the HomeCage Rack system overnight. After acclimation, behaviors were recorded over a 12-hour period. For analysis, the behaviors were grouped into 5 main categories: sleep, exploration, eating, drinking, and grooming, with specific behaviors assigned to these overarching categories. HomeCageScan was conducted at baseline and at 5 weeks post-intervention. Behavior data were recorded and analyzed for both duration (seconds) and frequency (bouts).^28^

### 2.7. Immunohistochemistry

Formalin-fixed, paraffin-embedded tissue sections (4 μm) were dewaxed in xylene and rehydrated through graded alcohols to water. Sections were blocked using a blocking buffer consisting of 1% bovine serum albumin, 0.1% cold fish skin gelatin (Biotium, Fremont, CA), 0.5% Triton X100 (Sigma Chemicals, St. Louis, MO), and 0.01% phosphate buffered saline (PBS, Sigma Chemicals, St. Louis, MO) for 30 min at room temperature. Slides were incubated with primary antibody diluted in the blocking buffer described above for 30 minutes at 37°C. The following antibodies were used in this study: calcitonin gene related protein (CGRP, 1:200 dilution in blocking buffer, Invitrogen, PAS-114929), and β-tubulin 3 (1:250, Invitrogen, 14-4510-82). Alexa 488nm conjugated goat anti rabbit IgG (H+L) (1:1000, Invitrogen, A32731) and Alexa 546nm goat anti rabbit IgG (H+L) (1:1000, Invitrogen, A11011) rabbit secondary antibodies respectively were diluted in the blocking buffer described above and applied for 30 minutes at 37°C. After the final wash step, slides were mounted in aqueous mounting media and stored for further analysis. For analysis images were captured and analyzed with a fluorescent Nikon Eclipse Ti2 microscope using NIKON Element software at 10× magnification (Nikon Inc, Melville, NY).

### 2.8 Statistical analysis

Three separate cohorts were used in this study: One cohort consisted of 10 male rats, the second cohort included 12 female rats, and the last cohort included 12 rats (6 male and 6 female). Rats in each cohort were divided into an amputation only group that consisted of a total of 15 rats (7 males, 8 females), and an amputation with iTMR cohort that consisted of a total of 19 rats (9 males, 10 females). Normality was assessed visually through QQ plots for each dataset, and all data met the assumption of normality. For comparisons between two groups, an unpaired t-test was conducted for each behavioral assay and intervention at each time point. First male and female rats were compared for sex-related differences with t test and segregated. After confirming normality, a two-way repeated measures ANOVA with Sidak post-hoc analysis was performed. If there were significant differences (p<0.05), the groups were segregated by sex for the intervention analysis. If there was no sex-related difference, intervention effect was analyzed in aggregate. Data are presented as mean ± standard error of the mean (SEM). Statistical analyses were carried out using GraphPad Prism 7 software (GraphPad Software, La Jolla, CA).

## 3 Results

### 3.1 Gross Assessment of Hindlimb Amputation Model

The hindlimb amputation surgery was generally well-tolerated by the rats. In the postoperative period, one rat developed a seroma at the amputation site, it was promptly irrigated and drained without signs of infection and went on to heal fully. No mortality was observed as a complication of the amputation such hemorrhage, infection, or mutilation.

From the immediate post-operative period and throughout the study, the animals maintained regular activity levels and did not exhibit any autotomy behaviors. They continued to gain weight at rates comparable to uninjured rats, with minimal to no hindrance in eating and drinking. No complications arose during any of the subsequent surgeries following the amputation.

At the conclusion of the experiments, autopsies were performed, and the coaptation sites were harvested. All rats in the iTMR group exhibited coapted nerves in continuity with no gross neuromatous growth. In contrast, rats in the amputation-only group showed enlarged nerve ends at the distal ligated nerve stumps consistent with neuroma formation. (Fig 2 A).

**Figure 2:**
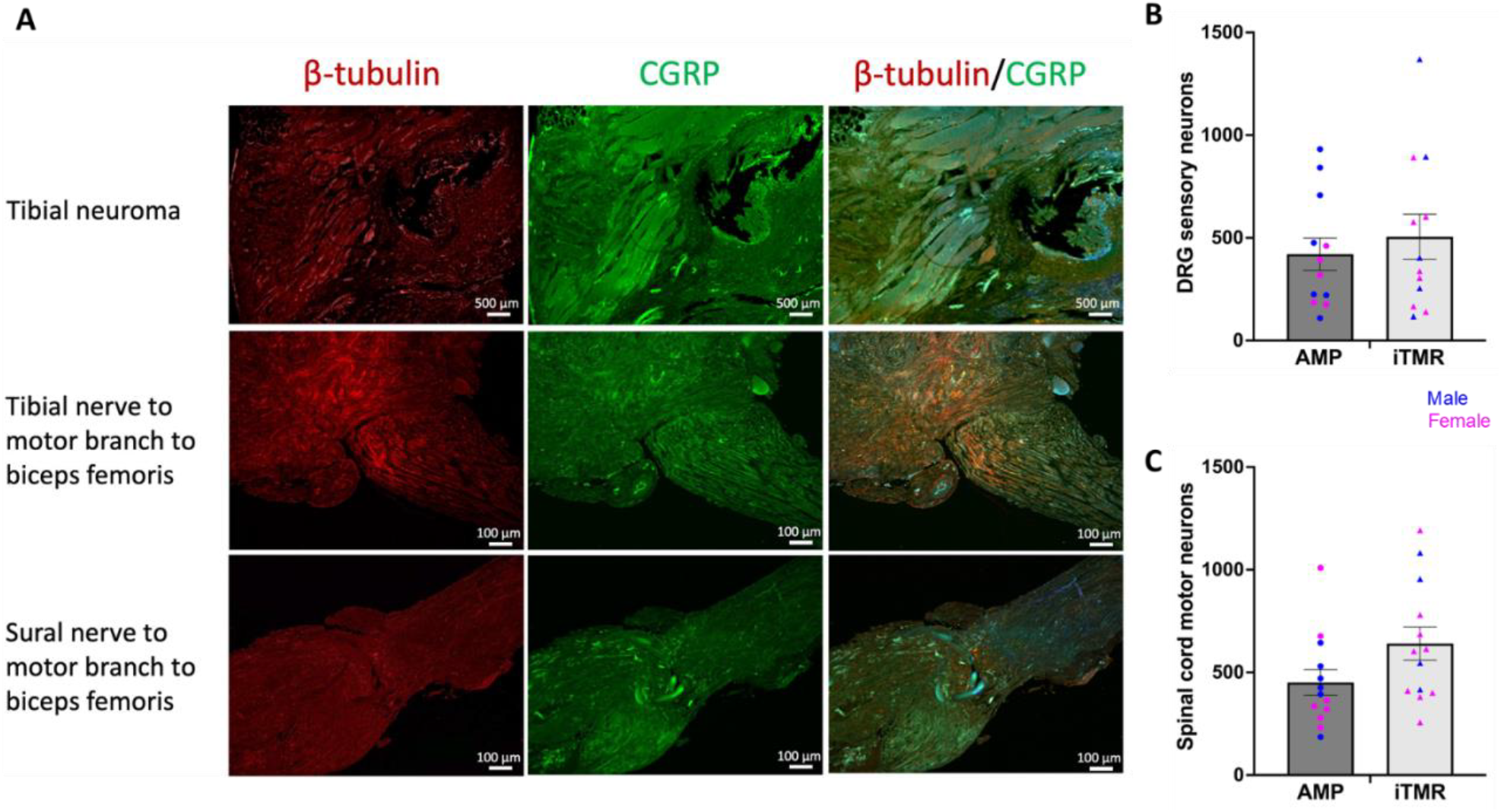
Comparison of Neuroma Formation and Neuronal Regeneration in AMP vs. iTMR Models. (A) Representative images comparing end neuromas to TMR coaptations. The tibial neuroma shows both sensory fibers (co-labeled with β-tubulin in red and CGRP in green) and a smaller proportion of motor fibers (β-tubulin staining only), presenting a disorganized structure. In contrast, the tibial and sural coaptations demonstrate no neuroma formation despite the size mismatch between donor and recipient nerves. Both sensory and motor fibers regenerate into the motor branch of the biceps femoris in the tibial nerve coaptation, while sensory fibers regenerate into the motor branch in the sural nerve coaptation. (B,C) Neuronal counts in DRG and spinal cord across AMP and iTMR cohorts. DRG sensory neuron and spinal cord motor neuron counts show no significant differences between AMP and iTMR groups (ns), indicating that iTMR does not affect the overall number of regenerating neurons. *p < 0.05, **p < 0.01. Sample sizes: n = 12-13 for amputation only (AMP) and n = 12-13 for amputation + iTMR (iTMR). Two-way repeated measures ANOVA with Sidak post-hoc analysis was performed for analysis.

### 3.2 Immunohistochemistry and characterization of retrograde labeling

Consistent with the gross examination, no neuroma formation was observed at the TMR coaptation sites. In contrast, at the distal end of the amputation-only nerves, a distended bulb containing disordered, predominantly CGRP-positive axons was present. Sensory fibers, identified by co-labeling with β-tubulin and CGRP, regenerated from the donor nerve to the recipient nerve (**Fig 2A**). Notably, the coaptation of the sural nerve to the motor branch of the biceps femoris, representing pure sensory-motor mismatch, demonstrated clear regeneration of sensory fibers into the smaller-diameter motor nerve.

The presence of iTMR did not affect the overall number of regenerating motor or sensory neurons. In amputation alone, 451±62 motor and 421±79 sensory neurons were preserved, and in iTMR, 641±80 motor and 505±110 sensory neurons were preserved **(Fig 2 B,C)**.

### 3.3 Intervention with TMR reduces hyperalgesia but not mechanical hypersensitivity

The pin test revealed that rats undergoing amputation with iTMR exhibited less hyperalgesia at 2 and 4 weeks post-amputation compared to the amputation-only group. At 2 weeks, there was a trend toward reduced hyperalgesia in iTMR rats (p = 0.09), with hyperalgesia responses of 26% ± 7 for the iTMR cohort and 45% ±8 for the amputation-only cohort. At 4 weeks, the difference became significant (p = 0.01), with hyperalgesia responses of 25% ± 5 for the iTMR cohort and 51% ± 8 for the amputation-only cohort (**Fig. 3A**). von Frey mechanical sensitivity testing revealed sex-related differences in the AMP group. Males exhibited significantly higher withdrawal thresholds compared to females at both 2 and 5 weeks post-intervention, indicating greater mechanical hypersensitivity in females. At 2 weeks, males had a mean withdrawal threshold of 6.72 ± 4.33, while females had 1.91 ± 0.85 (p = 0.01). At 5 weeks, this trend persisted, with males showing 6.72 ± 4.33 and females 1.91 ± 0.85 (p = 0.03) (**Fig 3B**)

**Figure 3:**
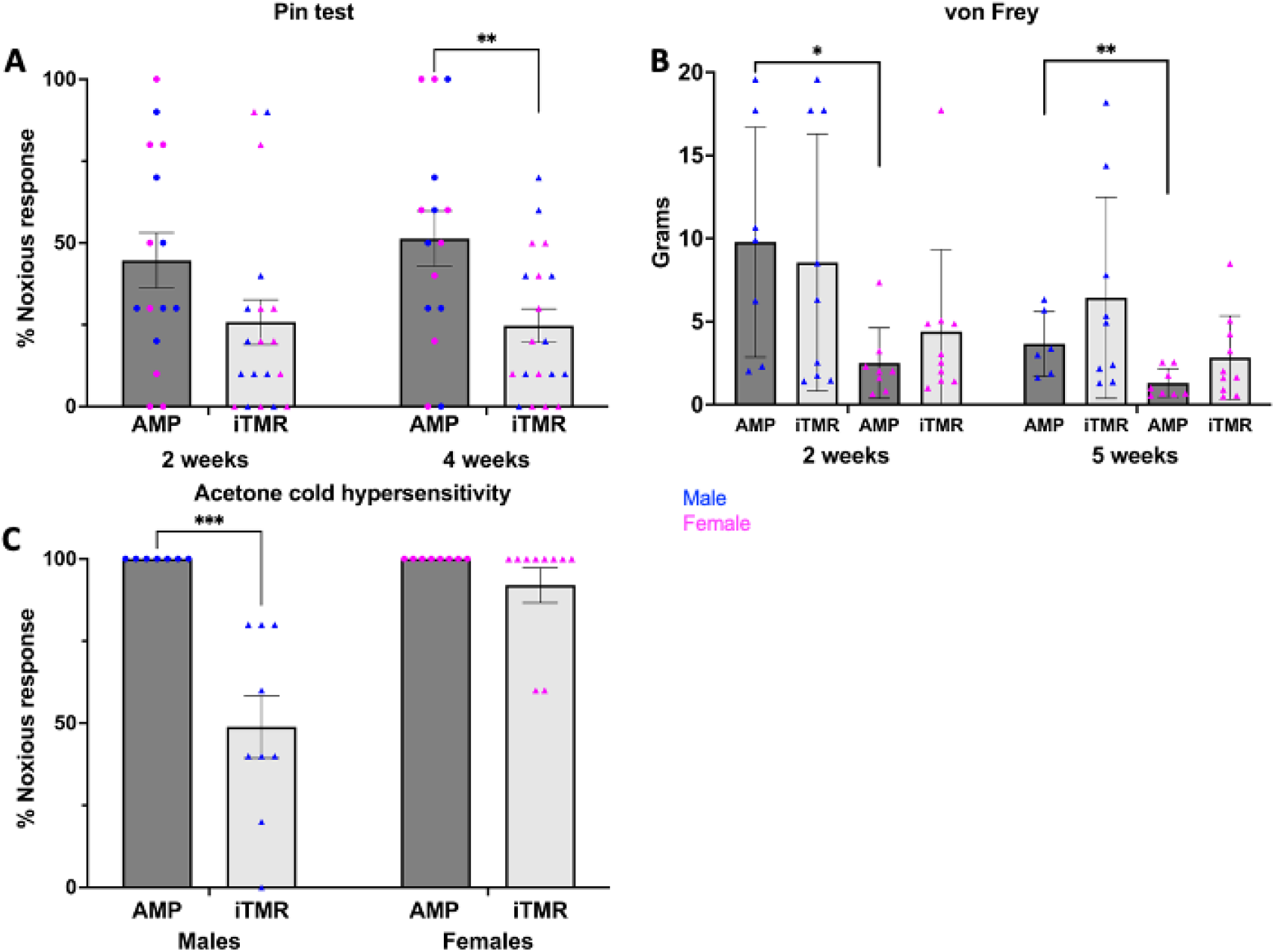
Sex Differences in Cold Hypersensitivity, von Frey Threshold, and Pin Testing Post-iTMR and Amputation-Only. (A) Pin testing showed that amputation with immediate TMR intervention reduced hyperalgesia at 2 and 4 weeks post-amputation compared to amputation-only. *p < 0.05, **p < 0.01. Sample sizes: n = 15-16 for amputation-only (AMP), n = 19 for amputation + iTMR. (B) von Frey mechanical sensitivity showed sex differences in the amputation-only group (AMP). Males had a significantly lower threshold compared to females at 2 and 5 weeks post-intervention, suggesting higher mechanical hypersensitivity in females. No significant differences were observed between males and females in the iTMR group. *p < 0.05, **p < 0.01. Sample sizes: n = 8-10 females, n = 7-9 males. (C) Acetone cold hypersensitivity showed a sex-specific response to intervention. Cold hypersensitivity 5 weeks post-intervention was significantly reduced in male amputees with iTMR compared to both amputation-only males and female amputees with iTMR. No significant differences were observed in cold hypersensitivity between females in either group. *p < 0.05, **p < 0.01. Sample sizes: n = 8-10 females, n = 7-8 males. Two-way repeated measures ANOVA with Sidak post-hoc analysis was performed for analysis.

### 3.4 Cold hypersensitivity was reduced with iTMR only in males

Cold hypersensitivity, measured 5 weeks post-intervention, demonstrated sexual dimorphism. Male amputees treated with iTMR exhibited reduced cold hypersensitivity compared to both male and female amputation-only subjects, and female amputees with iTMR. In contrast, all females developed cold hypersensitivity regardless of intervention (**Fig. 3C**).

### 3.5 Intervention with TMR reduces anhedonia

The iTMR cohort also showed a reduction in anhedonia, as evidenced by the sucrose preference test **(Fig. 4A)**. At 4 and 6 weeks post-intervention, these rats consumed significantly more sucrose compared to amputation-only counterparts that did not exhibit changes in sucrose consumption from baseline measurements taken prior to surgery. At baseline, there was no significant difference in sucrose preference between the groups (p = 0.15), with mean consumptions of 62 mL ±6 for the iTMR cohort and 49 mL ± 7 for the amputation-only cohort. At 4 weeks, iTMR rats consumed more sucrose than amputation-only rats (p <0.05), with means of 76 mL ± 9 and 43 mL ±5 respectively. At 6 weeks, iTMR rats consumed more sucrose than amputation-only rats (p <0.01) with means of 95 mL ± 11 and 58 mL ± 9 respectively. Further assessing whether there was a change from baseline between groups, AMP rats had no increase in sucrose consumed at either timepoint, but iTMR rats showed an increase in sucrose consumption from baseline (p<0.01).

**Figure 4:**
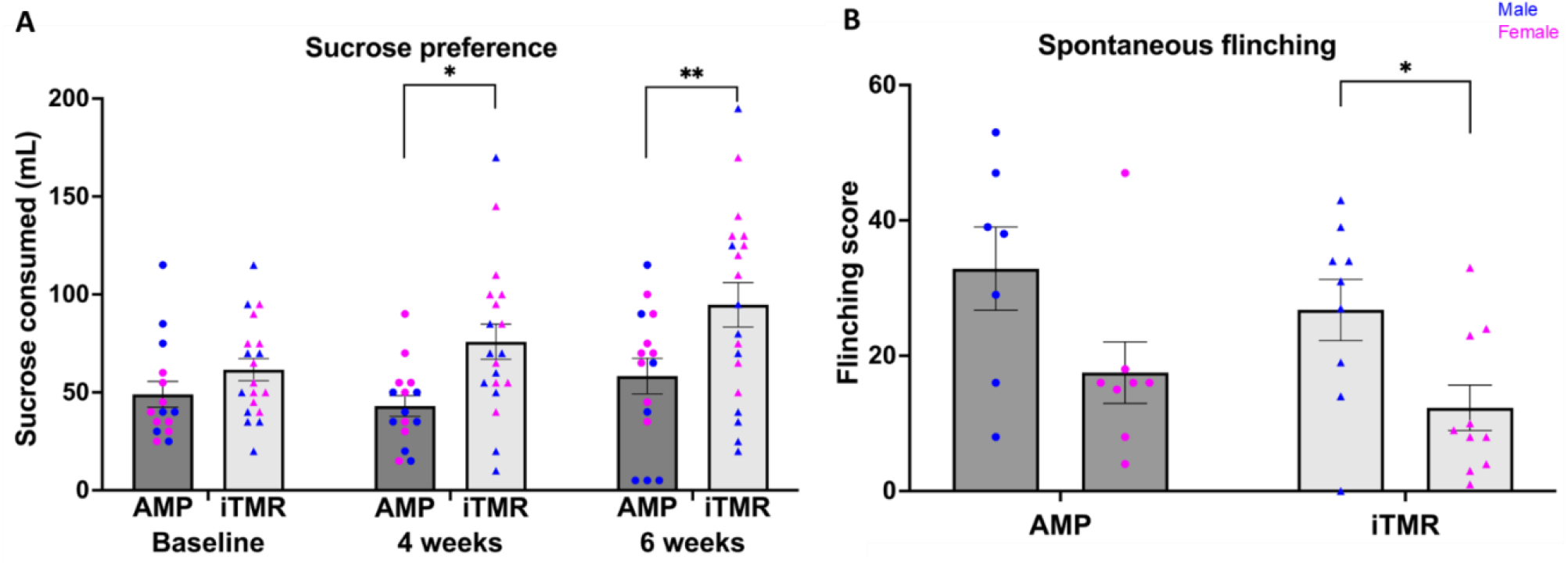
Reduced Anhedonia and Sex Differences in Spontaneous Pain Behaviors Post-iTMR. (A) Rats undergoing iTMR showed reduced anhedonia, as indicated by increased sucrose consumption at 4 and 6 weeks post-intervention compared to baseline. No significant changes were observed in sucrose consumption for amputation-only (AMP) rats. Baseline measurements were taken prior to surgery. *p < 0.05, **p < 0.01. Sample sizes: n = 14-15 for AMP, n = 19 for AMP + iTMR. (B) Spontaneous flinching behavior 5 weeks post-intervention. Males with amputation and iTMR exhibited significantly higher flinching scores than females in the same group, suggesting a sex-specific difference in pain-related behaviors. Flinching score represents the number of stump raises during a 2-minute observation period. *p < 0.05, **p < 0.01. Sample sizes: n = 8-10 females, n = 7-9 males. Two-way repeated measures ANOVA with Sidak post-hoc analysis was performed for analysis.

### 3.6 Males Exhibit More Spontaneous Pain Behaviors Than Females, Regardless of Intervention

Spontaneous flinching, an indicator of pain-related behavior, was assessed five weeks post-intervention. Male rats undergoing amputation with iTMR exhibited a higher number of spontaneous flinches compared to females with the same intervention (**Fig. 4B)**. Specifically, males with iTMR had a mean flinch score of 27 ±5 whereas females had a mean score of 12 ±3 (p <0.05). A similar trend was observed in the amputation-only cohort, in which females had a mean flinch score of 18± 5 and males had a mean score of 33± 6 (p = 0.06).

## Discussion

In this study, we developed a rat hindlimb amputation model and employed it to evaluate the analgesic effectiveness of iTMR. The model suitably replicates the clinical situation, as all rats are able to ambulate and maintain basic functions post-amputation, with minimal complications. Testing the stump for evoked pain behaviors shows that amputation results in mechanical hypersensitivity, hyperalgesia, and cold hypersensitivity. Intervention with iTMR reduces the severity of hyperalgesia and anhedonia in both males and females. Additionally, there is sexual dimorphism of iTMR effects as male rats, but not females, have reduced cold hypersensitivity when treated with iTMR.

### Pain Reduction and Behavioral Changes with iTMR

The magnitude of the evoked pain behaviors shown by the amputees were similar to those exhibited by rats undergoing the less severe injury of SNI. For example, the SNI-only cohort averaged ∼65% noxious response for the pin test and likewise, the amputees showed ∼48% noxious responses. Cold hypersensitivity noxious response were ∼83% for SNI and near 100% for amputees. ^12^ However, after amputation, the analgesia resulting from immediate TMR amputation is less complete, generally a reduction rather than a return to baseline. This may relate to the greater severity of the overall injury to the animal but may also reflect a limitation of testing the stump rather than the foot. An additional limitation posed by testing the stump is that baseline measures obtained from the foot are not entirely relevant as a baseline for the eventual stump. In the amputation model, the differences in pain behaviors between the amputation-only and iTMR groups emerged later than the differences observed between nerve injury and TMR groups in the SNI model. This delay may be attributed to the greater extent of tissue trauma in the amputation-including injury to muscle, bone, and skin-along with associated inflammation and edema. Overall, these findings align with clinical observations in human patients, where TMR, while beneficial, does not always eliminate pain.^7,9,29^

### Sexual Dimorphism in Pain Responses

Our study shows that male rats exhibit a reduction in cold hypersensitivity following iTMR, whereas females do not. Similar sex dimorphic analgesia findings were shown in the use of a regenerative peripheral nerve interface for nerve injury pain in rats.^30^ This work also demonstrated macrophage proliferation in the dorsal root ganglia (DRG) of treated male rats that was not observed in female rats, highlighting an immunomodulatory difference between sexes. Boullon et al. likewise showed increased cold allodynia in females but not male rats in the spared nerve injury model and observed an influence of the estrous cycle.^31^ In contrast, other neuropathic pain conditions and models have demonstrated sexually dimorphic effects on mechanical hypersensitivity and allodynia that we did not observe,^32^ suggesting the critical interplay of injury type and sex. These findings are important as much of the clinical studies of amputation related pain have assessed predominantly male populations.^7,9,33^ As TMR progresses to being widely available and standard of care, examining possible differential effects on male and female patients will be important.

### Regeneration: Motor versus

Our findings reveal no differences in motor or sensory neuron survival between iTMR and AMP groups, with neuron counts aligning closely with the normal range based on Swett et al.,^34^ who report a mean of 501 sensory neurons in the DRG, and a mean of 632 motor neurons. We do see that the standard deviation for the sensory neuron counts for the iTMR was double that of the amputation only group and this may be related to the relatively small size of the semimembranosus motor nerve branch and fragility in dissection for retrograde labeling. It was expected that there would be increased motor neuron retention in the iTMR group, for we observed a trend toward. TMR creates an environment favoring motor over sensory regeneration due to the modality mismatch in sensory-motor coaptation.^35-37^ Bergmeister et al.^38^ showed that when a much larger mixed motor-sensory nerve (representing 280 neurons) was coapted to a smaller motor branch (representing only 29 motor neurons) the retrograde labeled muscle was innervated by 56 neurons. Similarly, Lu et al.^39^ identified enhanced preservation of motor neurons when they transferred the cut end of the tibial nerve into the gastrocnemius in a procedure similar to TMR, although still a reduction compared to the native population. The implication of motor neuron retention or even competition with sensory axons for reinnervation through the coapted motor branch is not clear for pain outcomes. As the original purpose of TMR is to provide an EMG signal for a myoprosthesis, reinnervation with 30% of the native neurons is more than sufficient. As signal filtering continues to improve, this requirement is likely to be further minimized.^40^

### Limitations of the Study

Rat pain models, while valuable, cannot fully replicate the complexities of human pain. Neuron retention in rats may be greater than in humans due to a more robust axon sprouting and nerve regenerative capacity.^41^ Innervation of the skin at the created stump is not comparable to the plantar skin. However, this should not affect comparisons between treatment groups. Evoked behaviors are largely reflexive and do not measure the critical aspects of unpleasantness that is central to the definition of pain. However, the inclusion of non-evoked pain assessments, such as spontaneous guarding, flinching, and sucrose preference, provides additional insight into the animals’ affective state and pain experience. Furthermore, the consistency of withdrawal behaviors and licking of the stump remains reassuring in assessing reflexive pain responses.

### Future Directions

The rat hindlimb amputation model offers a new tool for exploring the effects and underlying mechanisms of nerve injury combined with limb loss. It may be used to model underrepresented populations by including females and a range of subject ages. Our findings highlight the potential of iTMR to reduce pain behaviors, with distinct sex differences in response to intervention. Future studies may focus on assessing changes over extended periods and on subjects with coexisting diseases such as diabetes, and further elucidating the mechanisms underlying the observed sexual dimorphism in pain and regeneration. Our findings also provide valuable insights into how timing may affect the effectiveness of TMR in clinical settings.

