## Supplemental Figures for "Rat Hindlimb Amputation Model Shows Analgesia and Sexually Dimorphic Cold Hypersensitivity with Immediate Targeted Muscle Reinnervation"

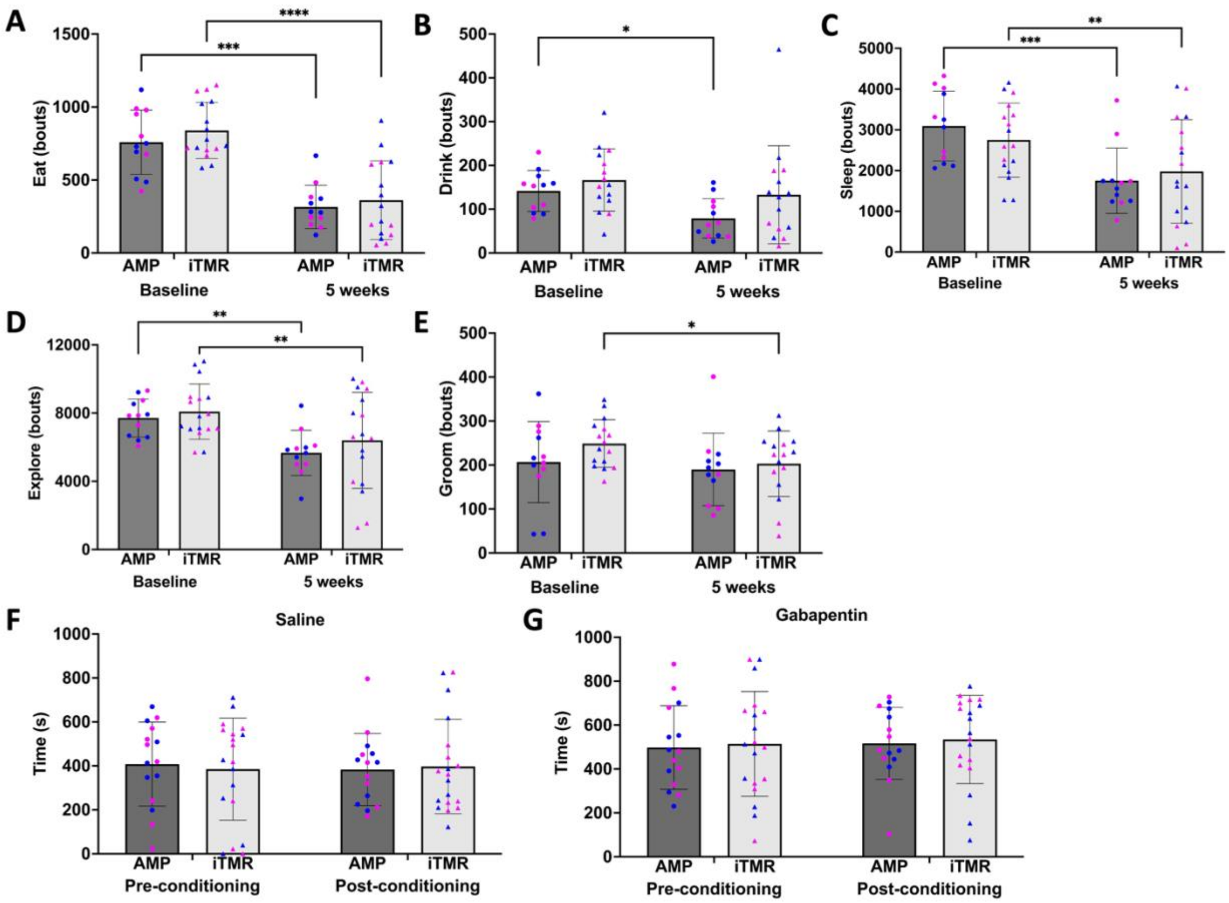


**Figure 5: HomeCage and Conditioned Place Preference Analysis with Gabapentin**

(A-E) Behavioral recordings from overnight HomeCage analysis showed no differences based on intervention or sex. Some time-based variations were observed, though these changes could not be linked to any specific pain phenotype. (F-G) In conditioned place preference (CPP) testing, neither cohort displayed a preference for the gabapentin-paired side. Contrary to expectations, rats with presumed higher pain phenotypes did not show increased time in the gabapentin-associated area during the post-conditioning phase.

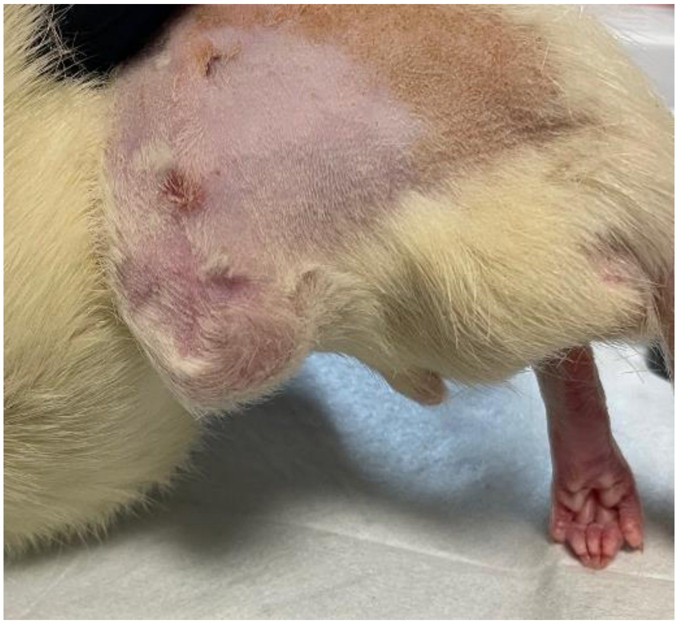

**Figure 6: Amputation**
